# Brain structural connectivity predicts brain functional complexity: DTI derived centrality accounts for variance in fractal properties of fMRI signal

**DOI:** 10.1101/826321

**Authors:** Josh Neudorf, Chelsea Ekstrand, Shaylyn Kress, Ron Borowsky

## Abstract

The complexity of brain activity has recently been investigated using the Hurst (H) exponent, which describes the extent to which functional magnetic resonance imaging (fMRI) blood oxygen-level dependent (BOLD) activity is self-similar vs. complex. For example, research has demonstrated that fMRI activity is more complex before than after consumption of alcohol and during task than resting state. The measurement of H in fMRI is a novel method that requires the investigation of additional factors contributing to complexity. Graph theory metrics of centrality can assess how centrally important to the brain network each region is, based on diffusion tensor imaging (DTI) counts of probabilistic white matter (WM) tracts. DTI derived centrality was hypothesized to account for the complexity of functional activity, based on the supposition that more sources of information to integrate should result in more complex activity. FMRI BOLD complexity as measured by H was associated with five brain region centrality measures: degree, eigenvector, PageRank, current flow betweenness, and current flow closeness centrality. Multiple regression analyses demonstrated that degree centrality was the most robust predictor of complexity, whereby greater centrality was associated with increased complexity (lower H). Regions known to be highly connected, including the thalamus and hippocampus, notably were among the highest in centrality and complexity. This research has led to a greater understanding of how brain region characteristics such as DTI centrality relate to the novel Hurst exponent approach for assessing brain activity complexity, and implications for future research that employ these measures are discussed.

## Introduction

Neuroimaging analyses have evolved since the advent of techniques such as functional magnetic resonance imaging (fMRI) to better understand what the blood oxygen level dependent (BOLD) signal implies about the underlying processing in the brain. For example, analyses have improved from a simple average of BOLD level during task vs. rest, to modeling expected activation time course based on an empirically derived hemodynamic response function convolved with the time course of stimuli presentations. Furthermore, resting state fMRI studies have revealed interesting baseline patterns of activation (i.e., resting state networks), and ushered in an age of network science in brain research. With the rise in resting state fMRI research, predicting resting state fMRI functional connectivity from diffusion tensor imaging (DTI) structural connectivity has been an important recent endeavor for researchers, and the use of graph theory analyses of structural networks have been integral to these investigations (e.g., Honey, Thivierge, & Sporns, 2010; Goñi et al., 2014; Bettinardi et al., 2017; Zhang et al., 2019; Chen, Hu, Chen, & Feng, 2019). Furthermore, there has been recent interest in predicting task-based fMRI activation from DTI structural connectivity (Osher et al., 2016; Ekstrand, Neudorf, Kress, & Borowsky, 2019). Beyond these fMRI activation descriptions of functional connectivity and task activation values, other novel methods to describe function include functional complexity, using metrics such as the Hurst exponent (H; e.g., Bullmore et al., 2001; Fadili, Bullmore, & Breatt, 2001; Shimizu, Barth, Windischberger, Moser, & Thurner, 2004). An important next step is to explore the extent to which one can predict the complexity in fMRI activation from the underlying DTI structure.

Since early demonstrations of using the Hurst exponent from the toolbox of chaos theory and fractals to analyse fMRI BOLD signal complexity (e.g., Bullmore et al., 2001; Fadili, Bullmore, & Breatt, 2001; Shimizu, Barth, Windischberger, Moser, & Thurner, 2004), interest has grown in this novel method for characterizing complexity of fMRI activation. The Hurst exponent has been applied to many domains of research including Alzheimer’s disease (e.g., Maxim et al., 2005), signal complexity change over the adult lifespan (Dong et al., 2018), distress related to medical treatment (Churchill et al., 2015), task versus rest states (He, 2011), and alcohol-induced intoxication versus non-intoxicated states (Weber, Soreni, & Noseworthy, 2014). Other measures of complexity such as functional variability have also demonstrated meaningful real-world associations (e.g., children develop more complex brain activity with age, see McIntosh, Kovacevic, & Itier, 2008). In the present research we apply network science graph theory metrics of *centrality* (i.e., how centrally important a particular region is to the network) as a novel way to further explore this important application of fractal analysis for characterizing fMRI activation complexity.

### Fractal Analysis of FMRI BOLD Signal Complexity

Fractal analysis of time course data measures the self-similarity or autocorrelation of the signal over different scales. High autocorrelation (or anti-correlation) signifies a “long-memory” process that is similar over shorter and longer time scales, whereas low autocorrelation signifies a “short-memory” process that varies between shorter and longer time scales. Complexity can be used as a descriptive analogous concept relating to these measures, whereby high autocorrelation is less complex than low autocorrelation, which is less complex than anti-correlation. The Hurst exponent (H) is one method of measuring signal complexity with fractal analysis, and describes the extent to which time course data such as fMRI BOLD signal activity is simple (H closer to 1) vs. complex (H closer to 0).

The complexity of brain activity has recently been investigated using H. Previous research has demonstrated that fMRI activity is more complex before consumption of alcohol than after consumption of alcohol (Weber, Soreni, & Noseworthy, 2014) and more complex during task than resting state (He, 2011). Aging has been associated with a reorganization of complexity, with left parietal and frontal regions losing complexity with age, while increases in complexity with age were observed in the insula, limbic system, and temporal lobe (Dong et al., 2018). Alzheimer’s disease has been associated with lower complexity than healthy controls in regions of the brain including medial and lateral temporal cortex, dorsal cingulate cortex, premotor cortex, left precentral gyrus, and left postcentral gyrus (Maxim et al., 2005). The measurement of H in resting state and task fMRI is a novel method that requires the investigation of additional factors contributing to BOLD signal complexity. One possible factor to be investigated is the connective structure of the brain network, which should play a vital role in determining the intrinsic patterns of coactivation of brain regions.

### Graph Theory Analysis of Brain Network Centrality

Graph theory analyses of networks such as the diffusion tensor imaging (DTI) structural connectivity network of the brain allow for assessment of how centrally important to the network a particular region is, via calculation of that region’s centrality. The most basic example of centrality is *degree centrality*, which is the count of connections between all other regions in the network to the region of interest. Other methods of calculating centrality represent more specific features of how centrally important to the network that regions is, such as weighting connections to the region based on how connected each other region is (*eigenvector centrality*; Bonacich, 1987), a method conceptually similar to eigenvector centrality that adjusts for the biased effect of largely connected regions on regions with low connections (*PageRank centrality*; Page, Brin, Motwani, & Winograd, 1999), counting the number of shortest paths between two regions in the network that pass through the region of interest (*betweenness centrality*; Anthonisse, 1971; Freeman, 1977), and average shortest path distance from the region to each other region in the network (*closeness centrality*; Bavelas, 1950). Variants based on current flow for betweenness centrality (*current flow betweenness centrality*; Brandes & Fleischer, 2005) and closeness centrality (*current flow closeness centrality*; Brandes & Fleischer, 2005; Stephenson & Zelen, 1989) capture similar features while using an electrical current model for information spreading, which may be more applicable to brain activity. Closeness and betweenness centrality are particularly sensitive to the topological role of a region in routing information between regions along shortest paths, and less sensitive to the number of connections. Graph theory centrality metrics allow for assessing specific ways that information is integrated and communicated through regions in the brain network (see Rubinov & Sporns, 2010 for a description of these methods).

### Relating Complexity to Centrality

Task vs. rest (He, 2011) and sober vs. intoxicated (Weber et al., 2014) states have been used to show that BOLD activity is more complex as measured by H during more organized cognitive states. Sufficient understanding has not yet been reached as to the reasons for increased complexity during these states, and one possibility is that this increased complexity during organized brain processing may be due to integrated co-processing of information from various brain regions concurrently. If this is the case, then regardless of task, regions that are more highly integrated via white matter (WM) tract connections to the rest of the brain network should have more complex activity than regions that are less connected to other brain regions, as these regions would be integrating information from more sources than other regions. The current research tests the hypothesis that DTI structural connectivity derived centrality measures sensitive to integration from many sources should be able to account for variation in the complexity of functional activity, based on the supposition that more sources of information to integrate should result in more complex activity.

## Materials and Methods

High quality DTI and resting state fMRI data for 100 unrelated subjects were obtained from the Human Connection Project (HCP) database (Van Essen et al., 2013; please see this paper for ethics statements). The HCP resting state fMRI data was used, which has been FSL *FIX* (Salimi-Khorshidi et al., 2014) preprocessed, white matter and cerebrospinal fluid activation regressed, and has undergone non-linear registration to Montreal Neurological Institute (MNI) standard space. All four sessions of this resting state fMRI data from HCP (4 sessions of 1200 acquisitions each, totalling 58 minutes) were used to calculate H for each voxel using the rescaled range estimation approach (e.g., first proposed by Hurst, 1951; see Hamed, 2007 for a discussion of rescaled range methods) in Python (Python Software Foundation, https://www.python.org) using adapted code from the *hurst* package (https://pypi.org/project/hurst/). The rescaled range estimation approach involves finding the rescaled range of partial samples of the time-series, having length *n*, defined as

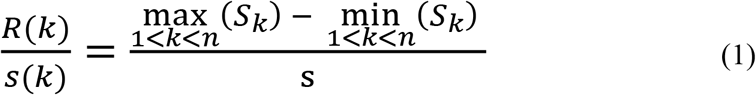

where S_k_ is the *k*th cumulative sum of deviations from the mean for each partial sample of varying sizes, which is given by

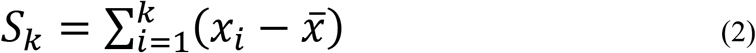

In equation 2, 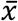 is the mean of the sample, and in equation 1, *s* is the standard deviation of the sample given by

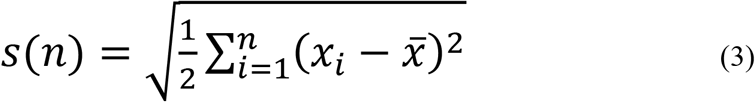

Finally, the Hurst exponent is estimated by finding the slope a least-squares regression line between log(R/s) and log(n) for each sample size n for which R/s was calculated, given that

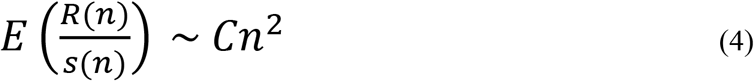

The mean H of the 4 sessions was calculated for each voxel, and then mean H was calculated for each of the 268 regions from the Shen functional atlas (Shen, Tokoglu, Papademetris, & Constable, 2013).

Probabilistic tractography was performed on the DTI data using FSL’s Graphics Processing Unit (GPU) implementation of *probtrackX* (Behrens et al., 2003; Hernandez-Fernandez et al., 2016; 2019) with a NVIDIA GTX 1080 graphics card, and the streamline pathway length correction option was applied. Importantly, the GPU implementation of *probtrackX* allowed for the computation of the 268 region connectivity matrix for more than one participant per day, allowing us to complete the calculations in approximately 3 months, whereas the default CPU implementation would have taken more than a week for each participant with the HCP high resolution DTI data (1.25mm × 1.25mm × 1.25mm resolution, 90 directions), which would have led to over 2 years of computation time. Probabilistic tractography is much more computationally intensive than deterministic tractography, and as such this resulting data represents an important contribution to the field of brain connectome research. The probabilistic tractography produced asymmetric structural connectivity matrices, which were converted to symmetric matrices by calculating the maximum streamline count between the cell in row *i* column *j*, and the cell in row *j* column *i*. The unthresholded connectivity matrix was then used to compute streamline weighted graph theory centrality measures of: 1) degree centrality (total connections to the region); 2) eigenvector centrality (number of connections weighted by degree of the connected region); 3) PageRank centrality (computes a ranking of the nodes based on the structure of the incoming links, originally designed as an algorithm to rank web pages; computed with α equal to the default of 0.85); 4) current flow closeness centrality (current flow method based on mean connectivity distance to other regions); and 5) current flow betweenness centrality (current flow method based on the number shortest paths the region is on). These graph theory metrics were calculated using the *NetworkX* python library, and the definitions provided are informed by the *NetworkX* documentation (Hagberg, Schult, & Swart, 2008; using functions *degree, eigenvector_centrality_numpy, pagerank, current_flow_closeness_centrality, and approximate_current_flow_betweenness_centrality*). The reciprocal of the DTI connectivity matrix was calculated and used as the weights for the current flow closeness centrality and current flow betweenness centrality, as these models determine shortest paths assuming that edge weights represent effective resistance in that path.

The means for H and degree centrality metrics were computed across subjects so that a mean value was assigned to each of the 268 brain regions. Mean H was then modeled as the criterion variable in a linear regression with centrality metrics as predictors. H was regressed on each of these centrality metrics separately and then multiple regression models were examined to investigate shared variance between the centrality metrics. Inter-individual correlation analyses were conducted to see if the effects were robust across regions, by examining for each region, when correlating centrality with H across all 100 subjects, how many regions out of 268 produced a significant positive or negative correlation, and whether the mean of this correlation is significant. Inter-region correlation analyses were conducted to see if the effects were robust across individual subjects, by examining for each subject, when correlating centrality with H across all 268 regions, how many subjects out of 100 produced a significant positive or negative correlation, and whether the mean of this correlation is significant. In order to assess inter-individual reliability, for each centrality metric and for H the values for each of the 268 regions for one subject are correlated with the 268 values for another subject, for all combinations of subjects (4950 combinations), which will result in a high positive correlation if there is good inter-individual reliability. The data that support the findings of this study are openly available in Zenodo at https://doi.org/10.5281/zenodo.3509709 (Neudorf, Ekstrand, Kress, & Borowsky, 2019).

## Results

Mean H was used as the criterion variable in simple and multiple linear regression models with centrality measures as predictor variables. For the simple regression models, degree centrality (*β* = −6.543 × 10^−3^, *t*(266) = −5.889, *p* < .0001, *R* = −.340; see Figure 1a), eigenvector centrality (*β* = −7.175 × 10^−3^, *t*(266) = −6.544, *p* < .0001, *R* = −.372; see Figure 1b), and PageRank centrality (*β* = −6.296 × 10^−3^, *t*(266) = −5.639, *p* < .0001, *R* = −.327; see Figure 1c) were found to be negatively associated with H, while current flow closeness centrality (*β* = 3.866× 10^−3^, *t*(266) = 3.341, *p* = .0001, *R* = .201; see Figure 1d), and log transformed current flow betweenness centrality (*β* = 3.481× 10^−3^, *t*(266) = 2.996, *p* = .0030, *R* = .181; see Figure 1e) were found to be positively associated with H (see Figure 1 for scatterplots of Hurst exponent as a function of centrality). The degrees of freedom used in these analyses correspond to the 268 regions of the brain, with each region averaged over the 100 subjects. In these linear regression models current flow closeness centrality and current flow betweenness centrality are positively associated with H, likely owing to their particular sensitivity to their position along shortest paths and distance from other regions, which measures routing rather than integration.

**Figure 1.**
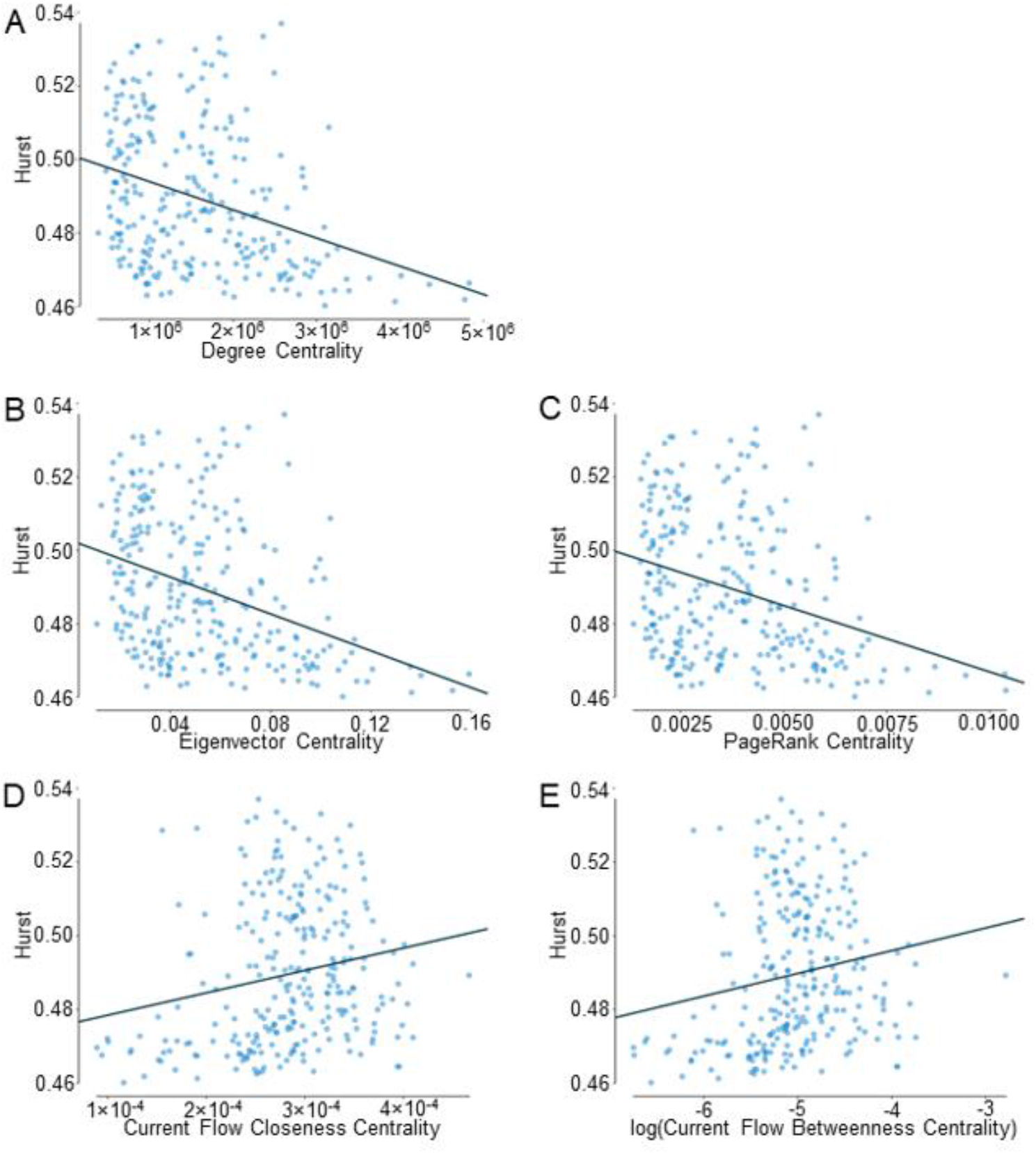
Scatterplots of Hurst exponent as a function of centrality, where centrality measures used are: a) degree centrality, *R* = −.340, *p* < .0001; b) eigenvector centrality, *R* = −.372, *p* < .0001; c) PageRank centrality, *R* = −.327, *p* < .0001; d) current flow closeness centrality, *R* = .201, *p* < .0001; and e) log(current flow betweenness centrality), *R* = .181, *p* = .0030.

Considering the high amount of collinearity between degree centrality, eigenvector centrality, and PageRank centrality, and also between current flow closeness centrality and log current flow betweenness centrality (see Table 1), these predictor variables were not included in the same multiple linear regression. A multiple linear regression of eigenvector centrality and current flow closeness centrality on H resulted in a significant model (*R* = .376, *R*^2^ = .141, *F*(2,265) = 21.78, *p* < .0001), with a significant negative effect of eigenvector centrality (*β* = −6.731× 10^−3^, *t*(266) = −5.581, *p* < .0001), and no significant effect of current flow closeness centrality (*β* = 1.065× 10^−3^, *t*(266) = .883, *p* = .3780). Including eigenvector centrality with log current flow betweenness centrality in a multiple linear regression model also resulted in a significant model (*R* = .376, *R*^*2*^ = .142, *F*(2,265 = 21.90), *p* < .001), with a significant negative effect of eigenvector centrality (*β* = −6.778× 10^−3^, *t*(266) = −5.807, *p* < .0001), and no significant effect of log current flow betweenness centrality (*β* = 1.156× 10^−3^, *t*(266) = .990, *p* = .3231). Analogous models including degree or PageRank centrality in place of eigenvector centrality yield the same pattern of results, but the models with eigenvector centrality consistently accounted for the most variance. These models demonstrated that the current flow centrality metrics do not account for a significant amount of variance when included in a multiple linear regression model with either degree centrality, eigenvector centrality, or PageRank centrality, and that the negative association between centrality and H is robust, whereby higher degree, eigenvector, and PageRank centrality is associated with lower H (higher complexity). Consistently in each of the degree centrality, eigenvector centrality, and PageRank centrality models, examples of the highest centrality regions, which also had some of the lowest values of H (highest complexity), included the left and right anterior and posterior regions of the thalamus (well-known to be important for connecting regions related to various different modalities), and regions of the medial temporal gyrus including the right posterior hippocampus (commonly viewed as being integral in the encoding of multi-modal information from many disparate brain regions, e.g., Damasio, 1989; see also Bird & Burgess, 2008 for a review; see Figure 2 for example sagittal and axial brain slices of degree centrality and H in various regions including the thalamus and hippocampus; see Table 2).

**Table 1.**
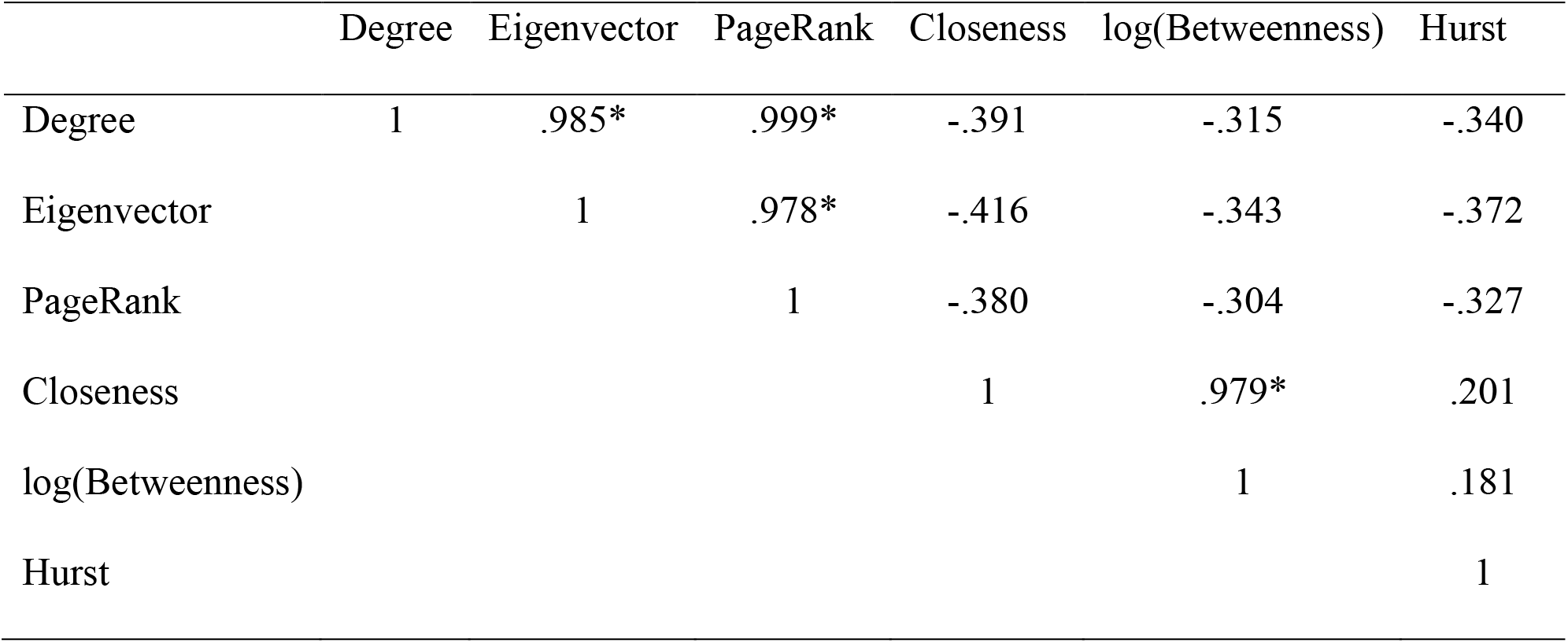
Predictor variable correlation matrix to assess multicollinearity. Values represent Pearson’s R correlation. Asterisks denote high collinearity (*R* > .95).

|  | Degree | Eigenvector | PageRank | Closeness | log(Betweenness) | Hurst |
| --- | --- | --- | --- | --- | --- | --- |
| Degree | 1 | .985* | .999* | -.391 | -.315 | -.340 |
| Eigenvector |  | 1 | .978* | -.416 | -.343 | -.372 |
| PageRank |  |  | 1 | -.380 | -.304 | -.327 |
| Closeness |  |  |  | 1 | .979* | .201 |
| log(Betweenness) |  |  |  |  | 1 | .181 |
| Hurst |  |  |  |  |  | 1 |

**Table 2.** Examples of highest degree, eigenvector, and PageRank centrality regions with some of the lowest H values.

| Region | Region Name | Eigenvector | Degree | PageRank | Hurst |
| --- | --- | --- | --- | --- | --- |
| 127 | Right Posterior Thalamus | .159 | $4.836 \times 10^6$ | .010 | .466 |
| 264 | Left Posterior Thalamus | .152 | $4.78 \times 10^6$ | .010 | .462 |
| 93 | Right Posterior Hippocampus | .140 | $4.35 \times 10^6$ | .009 | .466 |
| 263 | Left Anterior Thalamus | .136 | $3.94 \times 10^6$ | .009 | .461 |
| 128 | Right Anterior Thalamus | .136 | $4.01 \times 10^6$ | .009 | .468 |

**Figure 2.**
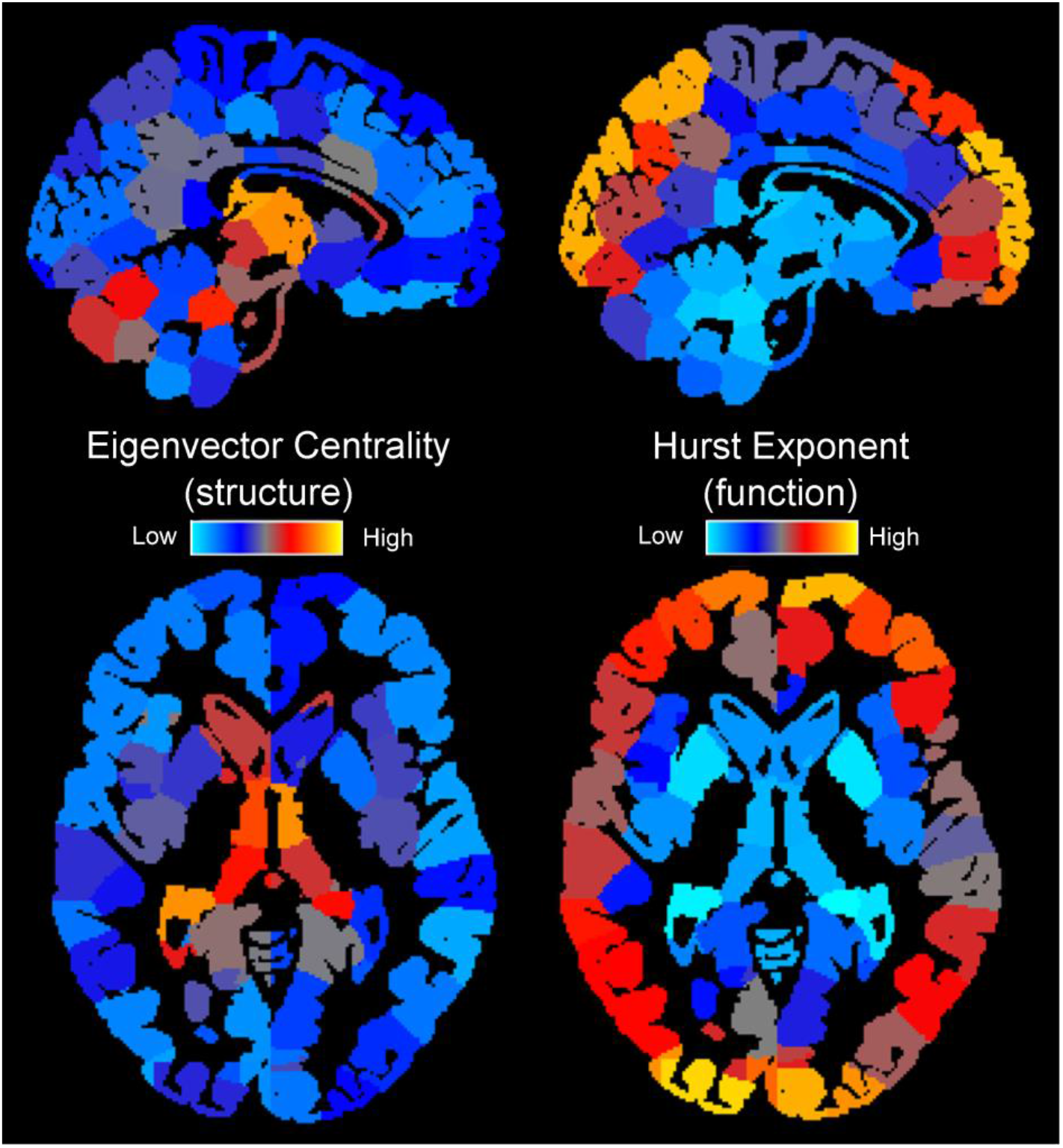
Example brain maps of degree centrality (left) and Hurst exponent (right), where yellow indicates higher values and blue indicates lower values. Centrality tended to be negatively associated with H, such that higher centrality (greater central importance to the network) was associated with lower H (more complexity).

### Inter-region Correlation Analysis

Given that the observed effect of centrality on H was relatively small (less than 20% of variance accounted for), additional analyses are conducted, first of all to see if this effect is robust across individuals. For all metrics, a large number of individual subjects had significant correlation values in the same direction as the primary analysis (negative for degree, eigenvector, and PageRank centrality; positive for current flow closeness centrality and log(current flow betweenness centrality)), and the mean correlation values for all metrics were also significant (see Table 3). Based on these additional analyses, we can say that the effects observed, especially for degree centrality, eigenvector centrality, and PageRank centrality, are robust across individuals.

**Table 3.** Inter-region correlation values between centrality metrics and H. Listed are the number of individual subjects (out of 100) showing significant positive or negative R values, the mean R value and SD, and the corresponding Fisher’s R-to-Z and p-value.

| Centrality Metric | # Subjects<br>sig. pos. R | # Subjects<br>sig. neg. R | R | SD | Fisher's R-to-Z | p |
| --- | --- | --- | --- | --- | --- | --- |
| Degree | 0 | 99 | -.290 | .062 | -4.853 | <.0001 |
| Eigenvector | 0 | 99 | -.303 | .069 | -5.088 | <.0001 |
| PageRank | 0 | 99 | -.282 | .061 | -4.723 | <.0001 |
| Closeness | 75 | 0 | .164 | .074 | 2.696 | .0070 |
| log(Betweenness) | 61 | 0 | .133 | .077 | 2.175 | .0296 |

*Note.* This analysis demonstrates that observed effects are robust across individuals.

### Inter-individual Correlation Analysis

In this case, none of the metrics produced consistent, significant correlation values (see Table 4). This analysis clearly demonstrates this effect is not driven by individual differences in the values of these metrics, but rather that the relative values of centrality and H are related within each individual’s brain, as seen in the inter-region correlation analysis.

**Table 4.** Inter-individual correlation values between centrality metrics and H. Listed are the number of regions (out of 268) showing significant positive or negative R values, the mean R value and SD, and the corresponding Fisher’s R-to-Z and p-value.

| Centrality Metric | # Subjects<br>sig. pos. R | # Subjects<br>sig. neg. R | R | SD | Fisher's R-to-Z | p |
| --- | --- | --- | --- | --- | --- | --- |
| Degree | 0 | 14 | -.086 | .076 | -.852 | .3944 |
| Eigenvector | 4 | 2 | .018 | .088 | .178 | .8590 |
| PageRank | 4 | 4 | .001 | .104 | .095 | .9243 |
| Closeness | 2 | 0 | .078 | .060 | .768 | .4423 |
| log(Betweenness) | 16 | 8 | .002 | .114 | .024 | .9805 |

*Note.* This analysis demonstrates that observed effects are not driven by individual differences in the values of these metrics.

### Inter-individual Reliability

All centrality metrics and H produced high reliability, with at least 4949 out of 4950 combinations of subjects producing a significant correlation, and high mean correlation values between .770 and .913 (see Table 5). This inter-individual reliability analysis indicates that the regional pattern of H and centrality metrics was highly reliable across individuals.

**Table 5.** Inter-individual reliability analysis values for centrality metrics and H. Listed are the number of subject combinations (out of 4950) showing significant positive R values, the mean R value and SD, and the corresponding Fisher’s R-to-Z and p-value.

| Variable | # Subjects<br>sig. pos. R | R | SD | Fisher's<br>R-to-Z | p |
| --- | --- | --- | --- | --- | --- |
| Degree Centrality | 4950 | .909 | .045 | 107.122 | <.0001 |
| Eigenvector Centrality | 4949 | .852 | .109 | 88.854 | <.0001 |
| PageRank Centrality | 4950 | .913 | .043 | 108.733 | <.0001 |
| Closeness Centrality | 4950 | .837 | .057 | 85.120 | <.0001 |
| log(Betweenness Centrality) | 4950 | .770 | .055 | 72.195 | <.0001 |
| Hurst Exponent | 4950 | .841 | .058 | 86.222 | <.0001 |

*Note.* The regional pattern of H and centrality metrics was highly reliable across individuals.

## Discussion

Fractal analysis of fMRI activity is a recent technique demonstrating that lower H is related to higher complexity of cognitive processing (e.g., Maxim et al., 2005; He, 2011; Weber et al., 2014). Our research demonstrated that higher complexity fMRI activity (lower H) can be explained in part by the centrality of the region in the structural DTI connectivity network. Specifically, degree centrality, eigenvector centrality, and PageRank centrality were negatively associated with H, meaning higher centrality on these measures was associated with more complex functional activity (lower H). Conversely, current flow closeness centrality and current flow betweenness centrality were positively associated with H, suggesting that these metrics are associated with less complex activation, which may be because they are particularly sensitive to simple routing of information rather than integration of information from many sources. Overall, in both simple and multiple regression analyses, eigenvector centrality consistently showed the strongest negative relationship with H (positive relationship with complexity), and as such may be the most robust metric of centrality to use in the future when associating DTI structural connectivity to resting state fMRI BOLD activity. Conversely, the interesting positive relationship with H demonstrated by current flow closeness centrality and current flow betweenness centrality was not robust, as it was not significant in the multiple regression model. The positive relationship between these current flow centrality metrics and H may be related to the sensitivity of these regions to topological location along shortest paths, which facilitate communication between distant regions in the network. In general, increased complexity with increased degree centrality suggests that the complexity of fMRI BOLD activation increases in the presence of integration of information from many sources.

With this evidence that complexity measured by H is related to centrality in the network, baseline expectations for complexity could be assessed at the level of each brain region using DTI structural connectivity networks before assessing task-based differences in fMRI complexity. Task fMRI data could also be examined with respect to complexity measured with H, as an indicator of information integration during tasks that manipulate integration of multiple modalities of information, such as semantic processing involving integration of action, shape, colour, emotional, and auditory modalities. Amodal semantic hubs hypothesized to be involved in integration of multiple modalities (e.g., the anterior temporal lobe; see Patterson, Nestor, & Rogers, 2007; Neudorf, Ekstrand, Kress, & Borowsky, 2019) may have higher complexity as measured by H than modal regions focused on a single modality (e.g., action, shape, colour, etc.) during semantic tasks, and other potential integrative hubs may also be identified in this way. Additionally, this knowledge of the relation of structural connectivity patterns to fMRI signal complexity could aid investigations into reorganization of brain signal complexity with age from the lens of comorbid structural connectivity reorganization (see Dong et al., 2018).

The investigation of the fractal indicator, H, of complexity and its relation to DTI structural network dynamics via graph theory centrality analyses undertaken in this research has demonstrated inherent differences in the complexity of fMRI BOLD signals can be accounted for by structural connectivity centrality of the various regions in the brain. This finding suggests that complexity is dependant not just on task (e.g., He, 2011) or on biological state (e.g., Weber et al., 2014), but also on the structural pattern of the connections as studied through graph theory analysis of DTI probabilistic streamlines. Supporting the validity of our approach, regions known to be highly connected, including the thalamus and hippocampus, were among the highest in centrality and complexity. The number of connections to other regions in the brain (degree centrality) is positively and consistently related to the complexity of brain activity as measured with fMRI, as are eigenvector and PageRank centrality. This research contributes to a greater understanding of the novel use of the Hurst exponent in assessing complexity of fMRI BOLD signals.

## REFERENCES

Anthonisse, J. M. (1971). The rush in a directed graph. (Technical Report BN 9/7). Stichting Mathematisch Centrum, Amsterdam.

Bavelas, A. (1950). Communication Patterns in Task-Oriented Groups. The Journal of the Acoustical Society of America, 22(6), 725–730. https://doi.org/10.1121/1.1906679

Behrens, T.E.J., Woolrich, M.W., Jenkinson, M., Johansen-Berg, H., Nunes, R.G., Clare, S., Matthews, P.M., Brady, J.M., & Smith, S.M. (2003) Characterization and propagation of uncertainty in diffusion-weighted MR imaging. Magn Reson Med, 50(5):1077–1088. https://doi.org/10.1002/mrm.10609

Bettinardi, R. G., Deco, G., Karlaftis, V. M., Van Hartevelt, T. J., Fernandes, H. M., Kourtzi, Z., … Zamora-López, G. (2017). How structure sculpts function: Unveiling the contribution of anatomical connectivity to the brain’s spontaneous correlation structure. Chaos: An Interdisciplinary Journal of Nonlinear Science, 27(4), 047409. https://doi.org/10.1063/1.4980099

Bird, C. M., & Burgess, N. (2008). The hippocampus and memory: insights from spatial processing. Nature Reviews Neuroscience, 9(3), 182–194. https://doi.org/10.1038/nrn2335

Bonacich, P. (1987). Power and Centrality: A Family of Measures. American Journal of Sociology, 92(5), 1170–1182. https://doi.org/10.1086/228631

Brandes, U., & Fleischer, D. (2005). Centrality Measures Based on Current Flow. In V. Diekert & B. Durand (Eds.), STACS 2005 (pp. 533–544). Springer Berlin Heidelberg.

Bullmore, E., Long, C., Suckling, J., Fadili, J., Calvert, G., Zelaya, F., … Brammer, M. (2001). Colored noise and computational inference in neurophysiological (fMRI) time series analysis: Resampling methods in time and wavelet domains. Human Brain Mapping, 12(2), 61–78. https://doi.org/10.1002/1097-0193(200102)12:2<61::AID-HBM1004>3.0.CO;2-W

Chen, Z., Hu, X., Chen, Q., & Feng, T. (2019). Altered structural and functional brain network overall organization predict human intertemporal decision-making. Human Brain Mapping, 40(1), 306–328. https://doi.org/10.1002/hbm.24374

Churchill, N. W., Cimprich, B., Askren, M. K., Reuter‐Lorenz, P. A., Jung, M. S., Peltier, S., & Berman, M. G. (2015). Scale-free brain dynamics under physical and psychological distress: Pre-treatment effects in women diagnosed with breast cancer. Human Brain Mapping, 36(3), 1077–1092. https://doi.org/10.1002/hbm.22687

Damasio, A. R. (1989). The Brain Binds Entities and Events by Multiregional Activation from Convergence Zones. Neural Computation, 1(1), 123–132. https://doi.org/10.1162/neco.1989.1.1.123

Dong, J., Jing, B., Ma, X., Liu, H., Mo, X., & Li, H. (2018). Hurst Exponent Analysis of Resting-State fMRI Signal Complexity across the Adult Lifespan. Frontiers in Neuroscience, 12. https://doi.org/10.3389/fnins.2018.00034

Ekstrand, C., Neudorf, J., Kress, S., & Borowsky, R. (2019). Structural connectivity predicts cortical activation during lexical and sublexical reading. Under review in Neuroimage.

Goñi, J., van den Heuvel, M. P., Avena-Koenigsberger, A., Velez de Mendizabal, N., Betzel, R. F., Griffa, A., … Sporns, O. (2014). Resting-brain functional connectivity predicted by analytic measures of network communication. Proceedings of the National Academy of Sciences of the United States of America, 111(2), 833–838. https://doi.org/10.1073/pnas.1315529111

Hagberg, A., Schult, D., & Swart, P. (2008). Exploring network structure, dynamics, and function using NetworkX. In G. Varoquaux, T. Vaught, and J. Millman (Eds), Proceedings of the 7th Python in Science Conference (11–15). Pasadena, CA USA.

He, B. J. (2011). Scale-Free Properties of the Functional Magnetic Resonance Imaging Signal during Rest and Task. Journal of Neuroscience, 31(39), 13786–13795. https://doi.org/10.1523/JNEUROSCI.2111-11.2011

Hernandez-Fernandez, M., Reguly, I., Giles, M., Jbabdi, S., Smith, S., & Sotiropoulos, S. N. (2016). A fast and flexible toolbox for tracking brain connections in diffusion MRI datasets using GPUs. Presented at the 22nd Annual Meeting of the Organization for Human Brain Mapping (OHBM), Geneva, Switzerland.

Hernandez-Fernandez, M., Reguly, I., Jbabdi, S., Giles, M., Smith, S., & Sotiropoulos, S. N. (2019). Using GPUs to accelerate computational diffusion MRI: From microstructure estimation to tractography and connectomes. NeuroImage, 188, 598–615. https://doi.org/10.1016/j.neuroimage.2018.12.015

Honey, C. J., Thivierge, J.-P., & Sporns, O. (2010). Can structure predict function in the human brain? NeuroImage, 52(3), 766–776. https://doi.org/10.1016/j.neuroimage.2010.01.071

Hurst, H. E. (1951). Long-term storage capacity of reservoirs. Transactions of the American Society of Civil Engineers, 116, 770–799.

Fadili, M. J., Bullmore, E. T., & Breatt, M. (2001). Wavelet methods for characterising mono- and poly-fractal noise structures in shortish fMRI time series. NeuroImage, 13(6, Supplement), 116. https://doi.org/10.1016/S1053-8119(01)91459-4

Freeman, L. C. (1977). A set of measures of centrality based on betweenness. Sociometry, 40.35−41.

Hamed, K. H. (2007). Improved finite-sample Hurst exponent estimates using rescaled range analysis. Water Resources Research, 43(4). https://doi.org/10.1029/2006WR005111

Maxim, V., Şendur, L., Fadili, J., Suckling, J., Gould, R., Howard, R., & Bullmore, E. (2005). Fractional Gaussian noise, functional MRI and Alzheimer’s disease. NeuroImage, 25(1), 141–158. https://doi.org/10.1016/j.neuroimage.2004.10.044

McIntosh, A. R., Kovacevic, N., & Itier, R. J. (2008). Increased Brain Signal Variability Accompanies Lower Behavioral Variability in Development. PLOS Computational Biology, 4(7), e1000106. https://doi.org/10.1371/journal.pcbi.1000106

Neudorf, J., Ekstrand, C., Kress, S., & Borowsky, R. (2019). FMRI of shared-stream priming of lexical identification by object semantics along the ventral visual processing stream. Neuropsychologia, 133, 107185. https://doi.org/10.1016/j.neuropsychologia.2019.107185

Neudorf, J., Ekstrand, C., Kress, S., & Borowsky, R. (2019). Data for: DTI derived centrality predicts fMRI complexity as measured by fractal analysis [Data set]. Zenodo. https://doi.org/10.5281/zenodo.3708483

Osher, D. E., Saxe, R. R., Koldewyn, K., Gabrieli, J. D. E., Kanwisher, N., & Saygin, Z. M. (2016). Structural Connectivity Fingerprints Predict Cortical Selectivity for Multiple Visual Categories across Cortex. Cerebral Cortex, 26(4), 1668–1683. https://doi.org/10.1093/cercor/bhu303

Page, L., Brin, S., Motwani, R., & Winograd, T. (1999, November 11). The PageRank Citation Ranking: Bringing Order to the Web. [Techreport]. Retrieved September 23, 2019, from http://ilpubs.stanford.edu:8090/422/

Patterson, K., Nestor, P. J., & Rogers, T. T. (2007). Where do you know what you know? The representation of semantic knowledge in the human brain. Nature Reviews Neuroscience, 8(12), 976–987. https://doi.org/10.1038/nrn2277

Rubinov, M., & Sporns, O. (2010). Complex network measures of brain connectivity: Uses and interpretations. NeuroImage, 52(3), 1059–1069. https://doi.org/10.1016/j.neuroimage.2009.10.003

Salimi-Khorshidi, G., Douaud, G., Beckmann, C. F., Glasser, M. F., Griffanti, L., & Smith, S. M. (2014). Automatic Denoising of Functional MRI Data: Combining Independent Component Analysis and Hierarchical Fusion of Classifiers. NeuroImage, 90, 449–468. https://doi.org/10.1016/j.neuroimage.2013.11.046

Shen, X., Tokoglu, F., Papademetris, X., & Constable, R. T. (2013). Groupwise whole-brain parcellation from resting-state fMRI data for network node identification. NeuroImage, 82, 403–415. https://doi.org/10.1016/j.neuroimage.2013.05.081

Shimizu, Y., Barth, M., Windischberger, C., Moser, E., & Thurner, S. (2004). Wavelet-based multifractal analysis of fMRI time series. NeuroImage, 22(3), 1195–1202. https://doi.org/10.1016/j.neuroimage.2004.03.007

Stephenson, K., & Zelen, M. (1989). Rethinking centrality: Methods and examples. Social Networks, 11(1), 1–37. https://doi.org/10.1016/0378-8733(89)90016-6

Van Essen, D. C., Smith, S. M., Barch, D. M., Behrens, T. E. J., Yacoub, E., & Ugurbil, K. (2013). The WU-Minn Human Connectome Project: An overview. NeuroImage, 80, 62–79. https://doi.org/10.1016/j.neuroimage.2013.05.041

Weber, A. M., Soreni, N., & Noseworthy, M. D. (2014). A preliminary study on the effects of acute ethanol ingestion on default mode network and temporal fractal properties of the brain. Magnetic Resonance Materials in Physics, Biology and Medicine, 27(4), 291–301. https://doi.org/10.1007/s10334-013-0420-5

Zhang, R., Shao, R., Xu, G., Lu, W., Zheng, W., Miao, Q., … Lin, K. (2019). Aberrant brain structural–functional connectivity coupling in euthymic bipolar disorder. Human Brain Mapping, 40(12), 3452–3463. https://doi.org/10.1002/hbm.24608

